# Histone N-tails modulate sequence-specific positioning of nucleosomes

**DOI:** 10.1101/2023.11.30.569460

**Authors:** Tatiana Nikitina, Wilfried M. Guiblet, Feng Cui, Victor B. Zhurkin

**Affiliations:** National Cancer Institute, National Institutes of Health, Bethesda, MD 20892, US; Thomas H. Gosnell School of Life Sciences, Rochester Institute of Technology, Rochester, NY, USA

## Abstract

The precise mechanisms governing sequence-dependent positioning of nucleosomes on DNA remain unknown in detail. Existing algorithms, taking into account the sequence-dependent deformability of DNA and its interactions with the histone globular domains, predict rotational setting of only 65% of human nucleosomes mapped *in vivo*. To uncover novel factors responsible for the nucleosome positioning, we analyzed potential involvement of the histone N-tails in this process. To this aim, we reconstituted the H2A/H4 N-tailless nucleosomes on human BRCA1 DNA (∼100 kb) and compared their positions and sequences with those of the wild-type nucleosomes. In the case of H2A tailless nucleosomes, the AT content of DNA sequences is changed locally at superhelical location (SHL) ±4, while maintaining the same rotational setting as their wild-type counterparts. Conversely, the H4 tailless nucleosomes display widespread changes of the AT content near SHL ±1 and SHL ±2, where the H4 N-tails interact with DNA. Furthermore, a substantial number of H4 tailless nucleosomes exhibit rotational setting opposite to that of the wild-type nucleosomes. Thus, our findings strongly suggest that the histone N-tails are operative in selection of nucleosome positions, which may have wide-ranging implications for epigenetic modulation of chromatin states.

## Introduction

Organization of eukaryotic DNA in chromatin is essential for many cellular processes such as gene expression, DNA replication and repair (1). The basic repeating unit of chromatin is the nucleosome core particle (NCP) containing ∼145 bp fragment of DNA tightly wrapped around histone octamer (2). Packaging of DNA into nucleosomes is defined by numerous factors including the DNA bending anisotropy (3-5) and interactions with histones (5-7), both of which are sequence dependent. While ATP-dependent chromatin remodeling enzymes and transcription factors can also determine nucleosome positioning patterns (8), they are beyond the scope of the present study.

Histone interactions with DNA involve both the histone globular domains and the tails. For the histone domain-DNA interactions, previous studies have identified 14 highly conserved ‘sprocket’ arginine residues deeply buried in the DNA minor groove (5, 9, 10). We use this notation (5) because the spatial arrangement of these arginines in the minor groove of nucleosomal DNA resembles the teeth of a sprocket meshing with the holes in the links of a bicycle chain. As to the histone tail-DNA interactions, the tails (and especially the N-tails) contain numerous lysine and arginine residues that serve as a platform for histone posttranslational modifications (11). Epigenetic alterations of these residues can lead to changes in chromatin states (12). For example, acetylation of lysine residues neutralizes positive charges on histones, which weakens the interaction between histone N-tails with negatively charged DNA phosphate groups. As a result, a closed chromatin state can be transformed into an open state with increased DNA accessibility. Note, however, that there are no published data on the local repositioning of nucleosomes caused by histone modifications.

Sequence-dependent bending anisotropy dictates how DNA is wrapped around a histone octamer (3-5). Previous studies have established various DNA sequence patterns favorable for nucleosome formation (13-24), including the well-known WW/SS pattern (24), see Figure 1. (Here and below, W stands for A or T, and S is for G or C.) Specifically, the SS dinucleotides are enriched in the positions where DNA bends into the major groove, while the WW dinucleotides are predominantly located in the sites where nucleosomal DNA bends into the minor groove. The ‘sprocket’ arginines penetrating minor groove at these sites, interact favorably with AT-rich DNA having electro-negative minor groove (5). These energetically favorable arginine-DNA contacts that occur every 10-11 bp in NCPs, provide structural basis for the rotational positioning of the WW/SS nucleosomes.

**Figure 1.**
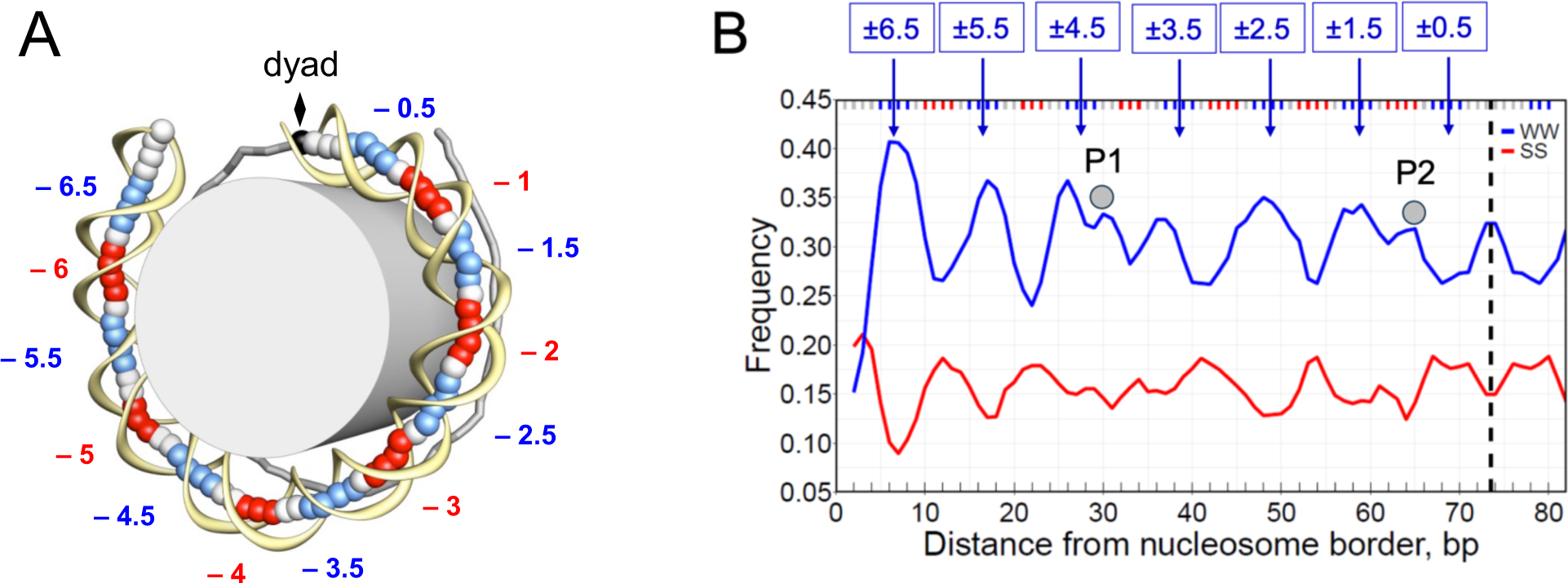
Locations of the minor- and major-groove bending sites and distributions of WW and SS dinucleotides along nucleosomal DNA. (A) The crystal structure of the 1kx5 nucleosome core particle with 147-bp long DNA (29) shown schematically: the DNA fragment is divided into two halves, separated by the dyad (diamond). The base-pair centers in the ‘ventral’ half are represented by large balls, and the sugar-phosphate backbone is shown by yellow ribbon. In the ‘dorsal’ half of nucleosome, sticks connect the base-pair centers. The minor- and major-groove bending sites are shown in blue and red, respectively. These sites are denoted according to their superhelical locations (SHL). (B) The frequencies of occurrences of combined AA, TT, AT, and TA dinucleotides (WW, shown in blue) and GG, CC, GC and CG dinucleotides (SS, shown in red) in each nucleosomal position, which were ‘symmetrized’ with respect to the dyad (dashed line). The three base-pair running averages of the WW and SS frequencies are presented for chicken nucleosomal DNA (24, 41). Gray circles indicate positions of the off-peaks, P1 and P2, near SHL ±4 and ±1.

In addition, the alternative DNA sequence pattern known as anti-WW/SS was described, in which the WW and SS profiles are in counterphase with the corresponding profiles observed in canonical WW/SS nucleosomes (25, 26). In this case, the ‘sprocket’ arginines seem to be in unfavorable contacts with GC-rich fragments in minor groove. Nevertheless, a recent study showed that the anti-WW/SS nucleosomes are widespread across different eukaryotes accounting for ∼25% in mammalian genomes (26). It remains unclear how these nucleosomes are stabilized.

We obtained a similar result earlier, when we compared several algorithms predicting positioning of human nucleosomes *in vivo* (27). Our algorithm (analyzing distribution of flexible pyrimidine-purine YR dimers as well as the WW/SS motifs) predicted 65% of the positions with 2-bp precision, compared to 55% for the widely used Kaplan-Segal model (28). In other words, 35-45% of nucleosome positions observed in vivo remain unaccounted for.

To tackle this problem, we compared the distributions of mono- and dinucleotides (A, T, W, WW and SS) along nucleosomal DNA across different eukaryotic genomes. In addition to the periodically oscillating sequence patterns observed earlier (24), we found two ‘rogue’ signals, one near superhelical locations (SHL) ±4 and the other near SHL ±1. Note that in published nucleosome structures (2, 29) the lysine-rich N-tails of the H2A and H4 histones are in favorable contact with the AT-rich DNA minor groove at SHL ±4 and SHL ±1/±2, respectively. Therefore, we hypothesized that the histone tails may play a role in the process of sequence-driven selection of the optimal nucleosome positions on DNA.

To elucidate a possible effect of the H2A and H4 N-tails on the sequence-specific formation of nucleosomes, we reconstituted the N-tailless NCPs and compared their positioning *in vitro* with that of the wild-type (W-T) NCPs. We found that overall, the H2A N-tailless NCPs are positioned in phase with the W-T NCPs (*i.e*., they are shifted by 10*n* bp, where *n* = 0, 1, 2, …). By contrast, a substantial number of the H4 N-tailless NCPs are in counterphase with the W-T NCPs (*i.e*., they are shifted by 10*n*+5 bp, like the anti-WW/SS nucleosomes are shifted *versus* canonical nucleosomes). Thus, our results suggest that preferential interactions of the histone N-tails with AT-rich minor groove may provide a novel mechanism for stabilization of anti-WW/SS nucleosomes observed *in vivo*. Implications for epigenetic control of nucleosome positioning are also discussed.

## Methods and Materials

### Nucleosome array reconstitution

Transformed *E. coli* with TAR/BAC containing full length hBRCA1 gene (BRCA1-TAR/YAC/BAC) was obtained from V. Larionov (30). It was first placed on agar plates supplemented with chloramphenicol as a selective agent (0.5 ml of 25 g/ml stock solution per 1 L of LB media). Cells were transferred into filtered LB media supplemented with 10 g yeast extract, 10 g NaCl, 2 g bactotryptone, 0.5 ml of 25 mg/ml of chloramphenicol and 0.4 ml of 5 M NaOH per 1 L of media and grown overnight at 37°C on platform shaker. Plasmids were prepared using Qiagen Large-construct kit (cat. # 12462) according to the manufacture’s instruction, and then cleaned with phenol chloroform extraction, precipitated, and resuspended in 10 mM Tris pH 8.0, 0.2 mM EDTA.

Reconstituted BRCA1-TAR/YAC/BAC arrays were assembled using *Xenopus* histone octamers with wild-type (W-T) and H2A/H4 N-tailless histones obtained from Histone Source, Protein Expression and Purification Facility at Colorado State University. Reconstitution was performed as described earlier (31). Briefly, core histone octamers and purified DNA were combined in a final mixture containing 2 M NaCl and 1 mM PMSF. Samples were kept on ice for 1 hour following dialysis in 3500 MWCO membranes against buffer (2 M NaCl, 10 mM Tris, 0.2 mM EDTA, 0.1% NP-40, 5 mM β-mercaptoethanol, pH=8.0) for 2 h. Samples were transferred to 1.5 M NaCl buffer and dialyzed for 2 h, followed by dialysis against 1 M NaCl and 0.75 M NaCl buffers, 3 hours each, and 0.5 M NaCl buffer overnight (all other buffer component were kept constant). Reconstitutes were next dialyzed against a buffer containing 5 mM NaCl, 10 mM Tris, 0.2 mM EDTA, 0.1% NP-40, 5 mM β-mercaptoethanol, pH 8.0, for 3 h. Samples were dialyzed in the buffer with 5 mM NaCl, 10 mM Tris, 0.025% NP-40 and no EDTA for additional 3 h.

Reconstituted chromatin (1 µg in 60 µl) in buffer containing 50 mM NaCl, 10 mM Tris 0.025% NP-40, 20% Ficoll, 0.7 mM CaCl_2_, and 2 mM MgCl_2_ was digested with 0.2 units MNase plus 5 units ExoIII for 10 min at 20°C. Reaction was stopped by addition of 20 mM EDTA and cooled on ice. Digested samples were treated with SDS and proteinase K for 2 h at 55°C and cleaned with Promega Wizard SV Gel and PCR Clean-Up System (cat. # A9282). Nucleosomal DNA fragments were subjected to paired-end sequencing on Illumina platform.

### In vitro nucleosome fragments

Raw FASTQ read pairs were trimmed with *cutadapt* (32). Trimmed read pairs were assembled with *PEAR* (33) to obtain nucleosome-size fragments. P-value for *PEAR* was set as *-p 0.0001*. The read pairs that failed to assemble with this p-value were discarded. The resulting fragments were then mapped with Bowtie2 (34, parameter *-x*), using the sequence of the *BRCA1* gene (from human genome hg19) as a reference. Fragments with *samtools* (35) flag 260 (unmapped/not primary alignment) were filtered out. To calculate mono- and di-nucleotide frequency profiles in nucleosomes, we selected DNA fragments with the lengths corresponding to the peaks of distributions presented in Figure 2: 147-149 bp for W-T, H2A and H4-del datasets, and 146-148 bp for H4-glob dataset.

**Figure 2.**
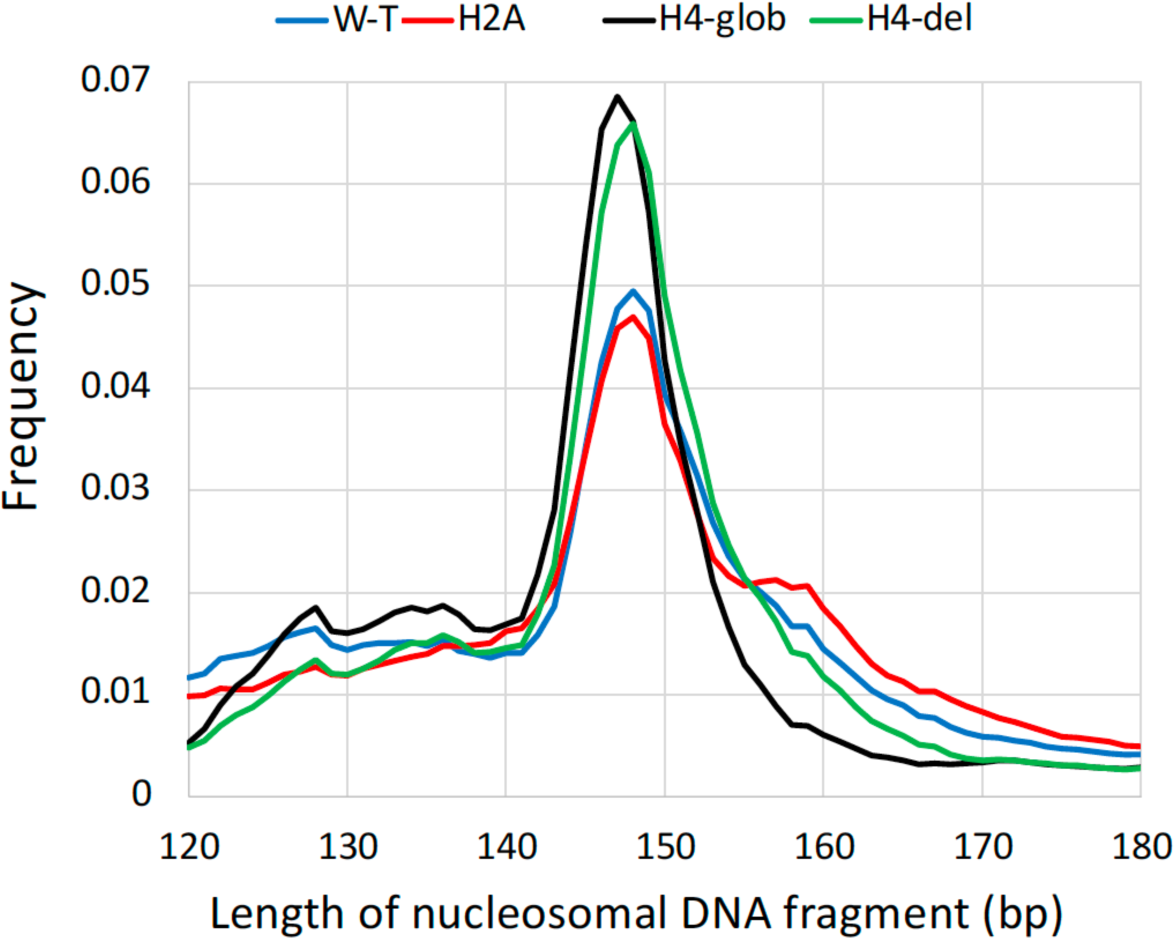
Distributions of the lengths of nucleosomes reconstituted with the wild type (W-T) and H2A/H4 N-tailless histones. To analyze distributions of mono- and di-nucleotides in nucleosomes, the DNA fragments with the following lengths were selected: 147-149 bp for W-T, H2A and H4-del datasets, and 146-148 bp for H4-glob dataset. The numbers of selected fragments were: 18 mil for W-T, 17 mil for H2A, 31 mil for H4-del and 27 mil for H4-glob datasets.

### In vivo nucleosome datasets

Several nucleosomal DNA datasets obtained *in vivo* were used in this study. Two datasets, one from yeast (36) and the other from mouse (37), were generated by the chemical method, and the dyad positions were precisely mapped at the single base-pair resolution. The other datasets were produced by paired-end MNase-seq from yeast (38), *Drosophila* S2 cells (39), mouse embryonic stem cells (37), and human lymphoblastoid cell lines 18508 and 19238 (40). BAM files were either downloaded from NCBI GEO database or obtained by mapping raw reads to the corresponding genomes using the default settings of Bowtie 2 (34), *i.e*., --sensitive, -I 0, -X 500. Only the fragments of 147 bp in length were used to calculate mono- and di-nucleotide frequency profiles.

## Results

### Periodically oscillating sequence motifs in nucleosomal DNA

The periodically oscillating nucleosomal DNA sequence motifs have been observed in the pioneer study by Satchwell, Drew and Travers (24) who analyzed distributions of all dimeric and trimeric steps in chicken nucleosomes *in vivo*. Their observation that the AT-rich and GC-rich motifs preferentially bend into the minor and major groove, respectively, was confirmed by numerous studies (41-43). These collective efforts have led to the widely accepted WW/SS scheme which we are using to represent the sequence-dependent deformability of nucleosomal DNA (Figure 1B). Note that the maxima of the WW profile occur in the minor-groove bending sites (SHL ±6.5, ±5.5, ±4.5, ±3.5, ±2.5, ±1.5), whereas the SS maxima occur in the major-groove bending sites (SHL ±6, ±5, ±4, ±3, ±2) (Figure 1B). Besides the periodically oscillating WW peaks, two pairs of distinctive ‘off-peaks’ P1 and P2 are clearly visible (Figure 1B), one near SHL ±4/±4.5 (P1) and the other at SHL ±1 (P2). Since no distortion in DNA trajectory is observed at these sites in nucleosome structures (2, 29), potentially, the irregularities in the WW profile may indicate the existence of certain histone-DNA interactions, in addition to those involving the ‘sprocket’ arginine residues (5-7, 9, 10).

To elucidate the sequence specificity of these off-peaks, we calculated the high-resolution profiles of mononucleotides A and T as well as the dinucleotide WW in nucleosomal DNA obtained from various species (see Figure 3, where the off-peaks P1 and P2 are marked by grey circles). Here we consider only 147-bp nucleosomal DNA fragments selected from millions of mapped NCPs, thereby increasing resolution of the sequence profiles, compared with the early studies (24). Note that in addition to the chicken MNase dataset (Figure 1B), the off-peak P1 is observed in the mouse chemical and the human MNase datasets (Figures 3D, 3F), while the off-peak P2 is present in all datasets. In most cases, it is detectable in the WW and A profiles, except the yeast MNase where it is clearly visible only in the A profile. Thus, we see that the off-peaks P1 and P2 are not an artefact observed only in a single species or after specific treatment (*e.g*., as a result of MNase or chemical cleavage); rather, they are inherent in a wide spectrum of eukaryotes.

**Figure 3.**
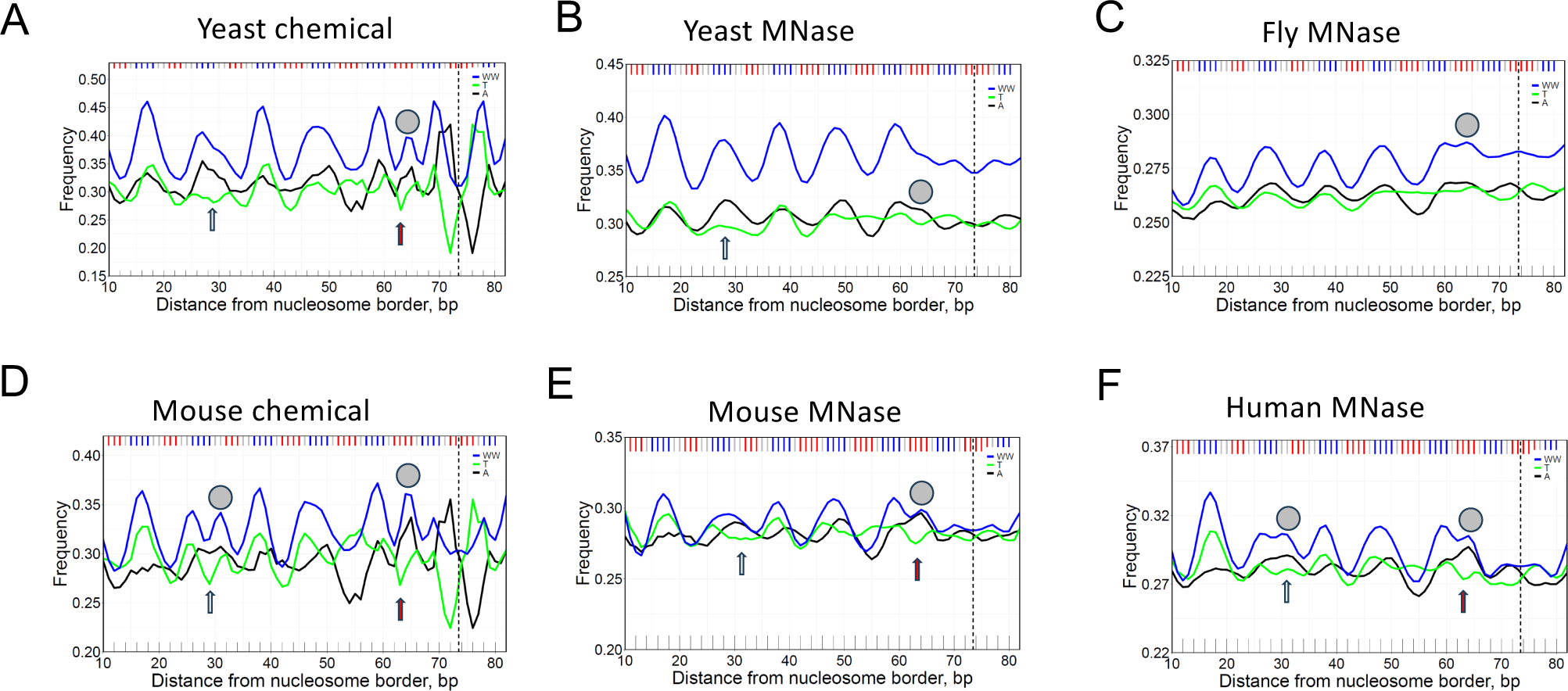
The frequencies of occurrences of nucleotides A and T as well as dinucleotide WW presented for yeast (A, B), fly (C), mouse (D, E) and human (F) nucleosomal DNA. The yeast and mouse nucleosomes are represented by two datasets, obtained with chemical mapping (A, D) and with MNase-seq mapping (B, E). The off-peaks, P1 and P2, are marked by gray circles (see Figure 1B). Arrows indicate the DNA regions with strand-specific preference A > T (SHL ±4, ±1). Note that at SHL ±1 the A > T effect is the strongest for mouse and human.

Both off-peaks P1 and P2 have a clear strand-specific preference for A *versus* T (see arrows in Figure 3). This effect was described by us earlier for yeast (43). The A > T preference at SHL ±4 (P1) was attributed to wedge formation by AA:TT dimeric steps observed in this region of alpha-satellite nucleosomal DNA crystal structure (29). According to this interpretation, the AA:TT wedges (15, 20) facilitate anisotropic DNA bending in nucleosome (4, 16), therefore, the AA dimers are more abundant in the leading strand at this location. This explanation is not universal because it does not account for the off-peak P2 at SHL ±1.

Note, however, that the locations SHL ±4 and SHL ±1 (where the off-peaks P1 and P2 are positioned) coincide with locations where the N-tails of H2A and H4 histones are in close contact with the DNA minor groove (Supplementary Figure S1). Since the nucleosomal DNA is relatively AT-rich (and depleted of the guanine amino-groups) at these locations, its minor groove is electro-negative and attractive for the positively charged histone N-tails. Hence, we posit that the histone N-tails may be responsible, at least partially, for formation of the off-peaks P1 and P2 (characterized by an increased occurrence of WW dimers).

### Histone H2A N-tailless constructs in vitro

To examine the role of histone N-tails in the selection of optimal nucleosomal DNA sequences *in vitro*, we compared the local AT content in nucleosomes reconstituted with the wild-type (W-T) and the H2A/H4 N-tailless histones (see Methods). For brevity, the N-tailless NCPs are denoted H2A, H4-glob and H4-del. First, we analyzed the combined frequencies of occurrences of adenines and thymines in the W-T nucleosomes and H2A constructs (Figure 4A) and found that both W(A+T) profiles exhibit periodic patterns with the peaks around the minor-groove bending sites, similar to the WW pattern of nucleosomes *in vivo* (Figures 1 and 3). Overall, the AT-content is lower by 0.4% in the W-T nucleosomes, except for the narrow interval near position #30 where the AT-content is higher by ∼2% in W-T.

**Figure 4.**
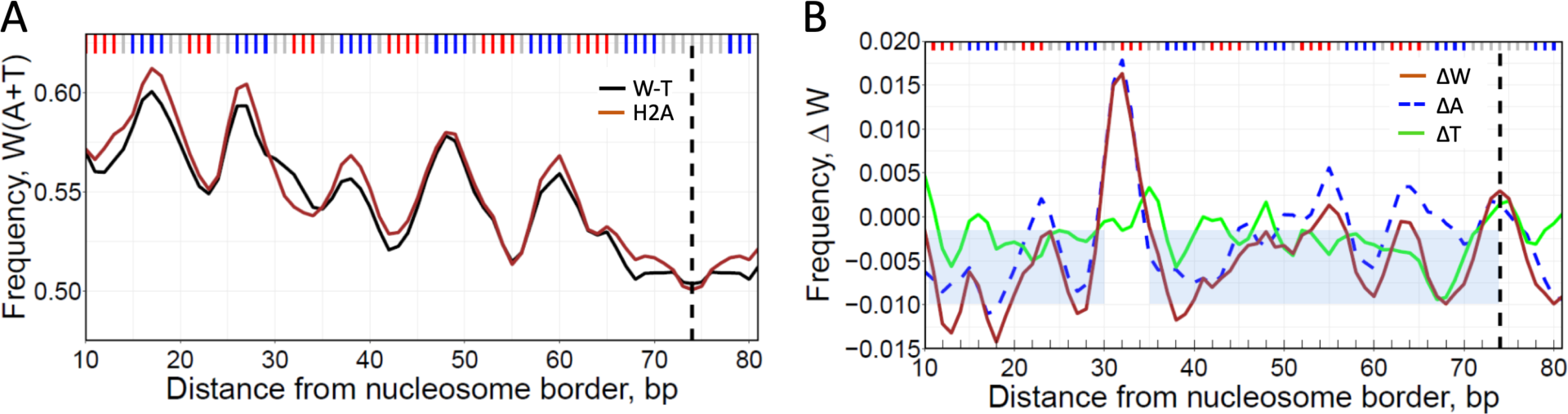
DNA sequence analysis of nucleosomes reconstituted with H2A N-tailless histones. (A) The frequencies of occurrence of W(A+T) nucleotides in the W-T (black) and H2A nucleosomes (brown). Three base-pair running averages of the frequencies were plotted over each nucleosomal position and symmetrized with respect to the nucleosome dyads (dashed lines). (B) ΔW, ΔA and ΔT profiles for H2A nucleosomes (brown, broken blue and green lines, respectively). The ΔW values were calculated as the W(A+T) frequency in W-T nucleosomes minus frequency in H2A nucleosomes for each nucleosomal position. The shaded areas represent the average ΔW = -0.006 and standard deviation of ΔW (0.004) in the ‘background’ regions 10-29 and 35-74.

To better visualize the changes in sequence composition between the two populations of NCPs, we calculated ΔW value representing the change in W(A+T) frequency upon transition from the N-tailless to the W-T nucleosomes (Figure 4B). The differential ΔW profile further demonstrates that the changes in W(A+T) are local, that is, the AT-content has increased noticeably in the region #30-34 (where H2A tail interacts with the DNA minor groove in X-ray structure (29), while in the other NCP regions the ‘compensatory’ changes in ΔW profile are relatively minor and exhibit elements of randomness (Figure 4B). Note that the ΔW value at position #32 is significantly higher than in the adjacent regions (p < 10^-43 in the one-sample t-test).

Importantly, the local increase in the AT-content occurs exclusively due to adenines (Figure 4B), which is entirely consistent with the strand-specific preference for A *versus* T described above *in vivo* (see Figure 3F, position #30). This result proves that indeed, modification of the histone H2A N-tails (in particular, their truncation) produces detectable changes in the W(A+T) frequency profile of the NCPs reconstituted *in vitro.* As shown below, these changes are accompanied by nucleosome repositioning (Supplementary Figure S2).

### Histone H4 N-tailless constructs in vitro

Removal of the H4 N-tail produced quite a different effect compared to H2A (Figure 5A). Instead of the local changes in W(A+T) frequency observed for the H2A construct, in the case of H4-del and H4-glob, the AT-content has decreased by 1-2% (relative to the W-T level) all over the length of a nucleosome. This is clearly visible in the differential ΔW profiles (Figure 5B). For the H4-del construct, ΔW exhibits strong peaks in the central part of nucleosome, at SHL 0 (dyad), SHL ±1 and ±2. For the H4-glob construct, the ΔW peaks are approximately two times higher and more widespread (Figure 5B). In other words, a complete truncation of N-tail (in H4-glob) leads to the more pronounced changes in the W(A+T) profile of nucleosome DNA sequences, compared to the partial removal of this tail (in H4-del).

**Figure 5.**
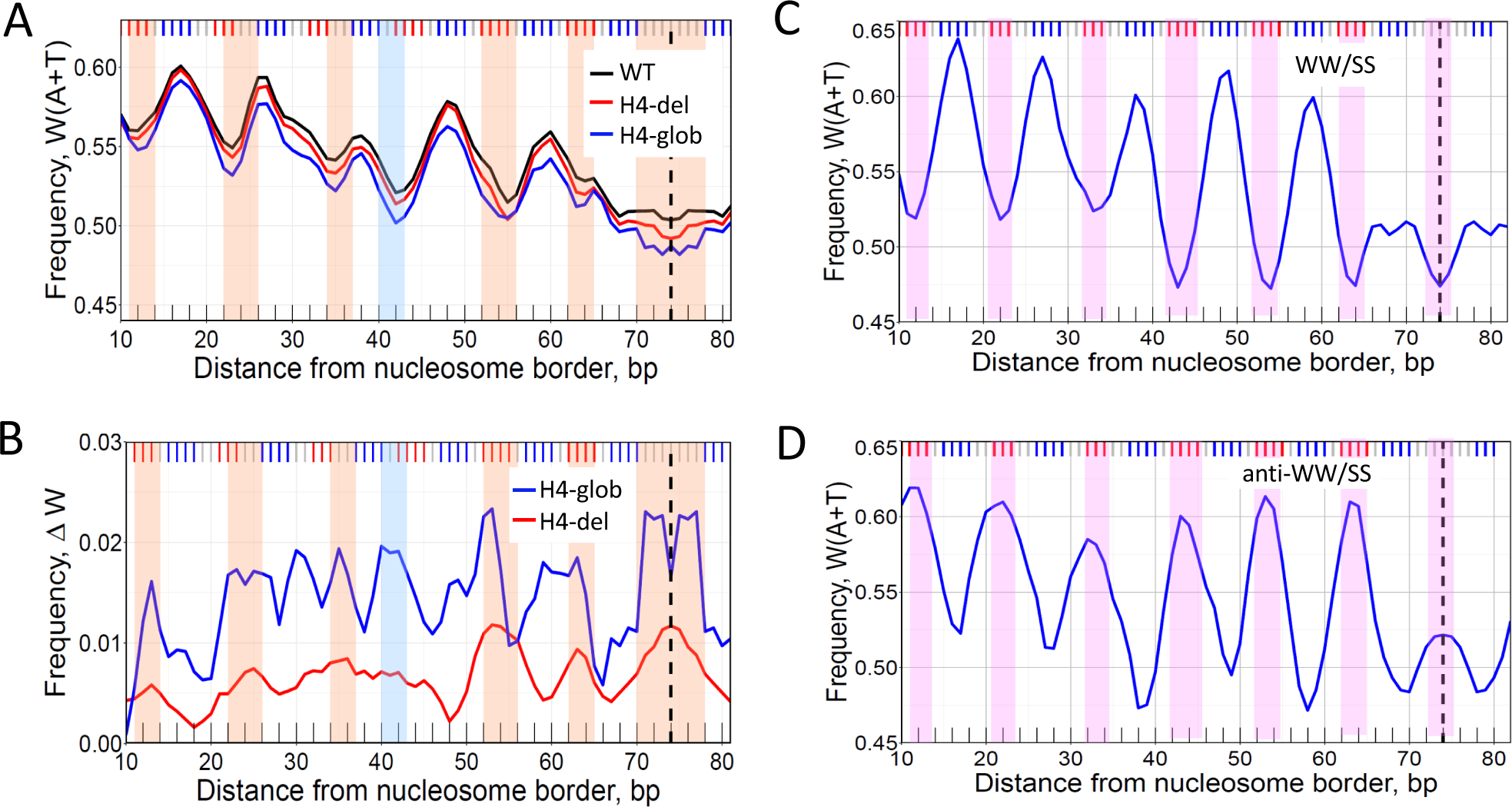
DNA sequence analysis of nucleosomes reconstituted with H4 N-tailless histones. (A) The frequencies of occurrence of W(A+T) nucleotides in the W-T nucleosomes (black), H4-del (red) and H4-glob (blue). (B) ΔW profiles for H4-del (red) and H4-glob (blue) nucleosomes. The ΔW values were calculated as in Figure 4. The orange vertical stripes emphasize close correspondence between the local minima in W(A+T) profile (A) and the local maxima in ΔW (B). The light blue vertical stripe indicates the local minima in W(A+T) at position #42 that have no corresponding local maximum in ΔW (H4-del). (C, D) W(A+T) profiles for the W-T nucleosomes having the canonical WW/SS pauern (C) and the ank-WW/SS pauern (D). See ref. (26) for descripkon of the detailed procedure. The two populakons of nucleosomes presented in (C, D) were denoted as Type 1 and Type 4, respectively (26). The pink vertical stripes indicate local minima in the W(A+T) profile for nucleosomes having the WW/SS pauern (C), and local maxima for nucleosomes having the ank-WW/SS pauern (D). Note that ank-WW/SS nucleosomes comprise 23% of all nucleosomes *in vitro*, in close agreement with the frackon of ∼25% for human nucleosomes *in vivo* (26).

Direct comparison of the ΔW and W(A+T) profiles shows that they are negatively correlated (Figures 5A, 5B). Indeed, each local minimum in W(A+T) corresponds to one of the local maxima in ΔW (H4-glob), as shown by vertical stripes in Figure 5. In the case of H4-del, six out of seven local minima in W(A+T) correspond to one of the local maxima in ΔW (H4-del). (The only exception is position #42; see the light blue vertical stripe.) In other words, the ΔW (H4) values are the highest where W(A+T) are the lowest. Thus, we observe two periodically oscillating signals in the nucleosomal DNA sequences – in addition to the canonical W(A+T) pattern, there is the ΔW pattern, associated with the H4 N-tail and positioned in counterphase with W(A+T).

In this regard, note that the ΔW (H4-glob) profile resembles the anti-WW/SS profile studied earlier (26) − both have peaks at the major-groove bending sites (Figures 5B, 5D), which is the opposite of the canonical W(A+T) profiles (Figures 5A, 5C). The rotational settings of the WW/SS and anti-WW/SS nucleosomes are counter-phased -- that is, the relative shift between these nucleosomes is ∼10*n*+5 bp (5, 15, 25, etc.). Therefore, one can expect that a similar shift of ∼10*n*+5 bp would be observed between the W-T and H4-tailless nucleosomes (Figures 5A, 5B).

### Inter-nucleosome distance cross-correla=on

To describe relative positions of nucleosomes in two populations we used the distance cross-correlation (DCC) function introduced earlier (42). For each pair of NCP positions (one from the first set, and the other from the second set), the distance ‘*dist*’ between the NCP dyads is calculated, and the occurrences of all distances are summed up (Figure 6). Note that multiple occurrences of nucleosomes in the same position are counted multiple times —that is, if the two NCP dyad positions are separated by 20 bp and occur 5 and 10 times, respectively, the corresponding DCC (*dist*=20) is calculated as 5 × 10 = 50.

**Figure 6.**
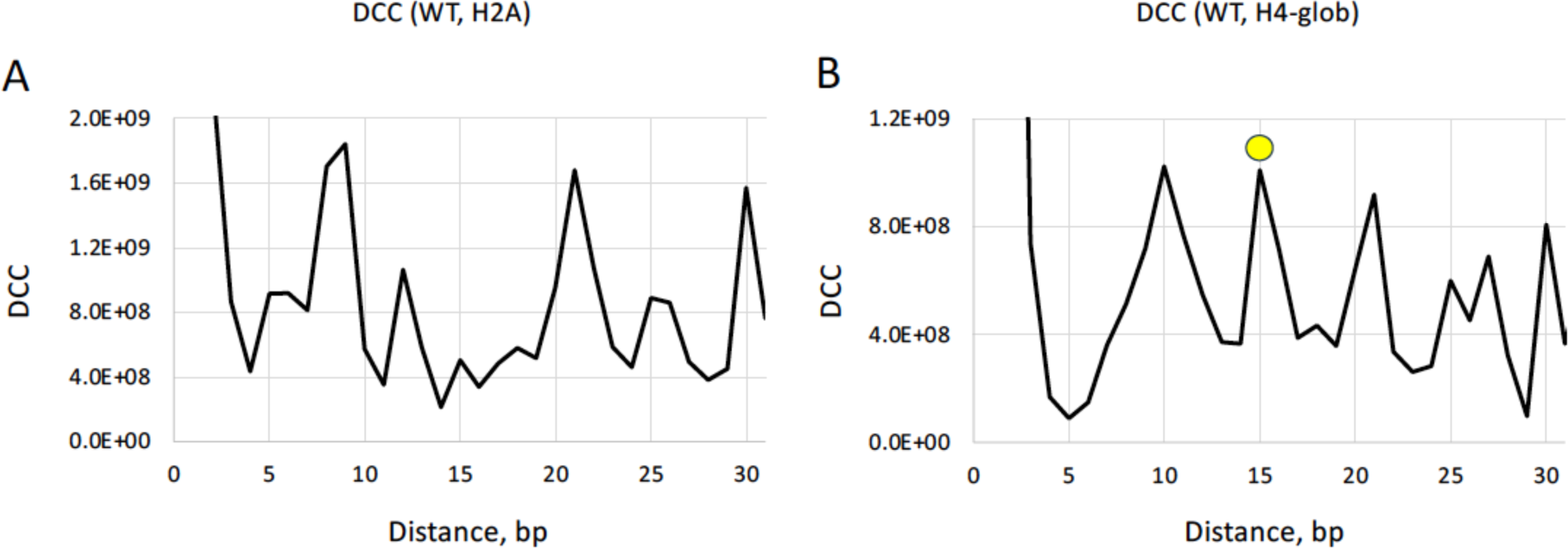
Inter-nucleosome distance cross-correlakon funckons (DCC). (**A**) Cross-correlakon between NCP posikons in the W-T and H2A datasets. (**B**) Cross-correlakon between NCP posikons in the W-T and H4-glob datasets. The peak at 15 bp is emphasized by yellow circle. The NCP posikons with occurrences > 5000 were included in calculakon of DCC (see Supplementary Figures S2, S3 for typical occurrence values).

A comparison of the W-T and H2A nucleosome subsets revealed strong DCC peaks at ∼10, ∼20 and 30 bp (Figure 6A), indicating that the NCPs from W-T and H2A are mostly separated by an integral number of DNA helical turns and represent the rotationally related translational positions (42, 44); see Supplementary Figure S2 where the W-T and H2A nucleosomes are shi∼ed by 20 bp.

In the case of W-T and H4-glob nucleosomes, the DCC profile has an additional peak at 15 bp (Figure 6B), suggesting that there are quite a few occurrences when nucleosomes from the two subsets have opposite rotational seSngs. The representative example is given in Supplementary Figure S3, where the NCP occurrence profiles are shown for 1 kb region of BRCA1 gene. Note that both profiles have twin peaks separated by 15 bp. In the W-T set, the dominant NCP position corresponds to the ‘left’ peak, whereas in the H4-glob set, the dominant NCP position coincides with the ‘right’ peak (Supplementary Figures S3A, S3B). Furthermore, the H4-del nucleosomes follow the same trend as H4-glob (Supplementary Figure S3C), and H2A nucleosomes are positioned similarly to W-T (Supplementary Figure S3D). We consider this result as the proof of principle demonstrating that predominant NCP positions in the W-T and H4-tailless populations can be separated by 10*n* + 5 bp (that is, can have opposite rotational settings).

### A novel mechanism for stabilization of the anti-WW/SS nucleosomes

According to our model, a nucleosome is stabilized by two types of histone-DNA interactions (Figure 7). The first type involves the ‘sprocket’ arginines in histone globular domains penetrating the DNA minor groove at locations SHL ±0.5, ±1.5, … ±6.5 (3-5). As mentioned in Introduction, these locations are characterized by a narrowed minor groove and are usually enriched with the WW dinucleotides (Figure 1). The positively charged arginines thus favorably interact with the electro-negative minor groove facing in, thereby selecting (and stabilizing) nucleosomes with the canonical WW/SS sequence pattern.

**Figure 7.**
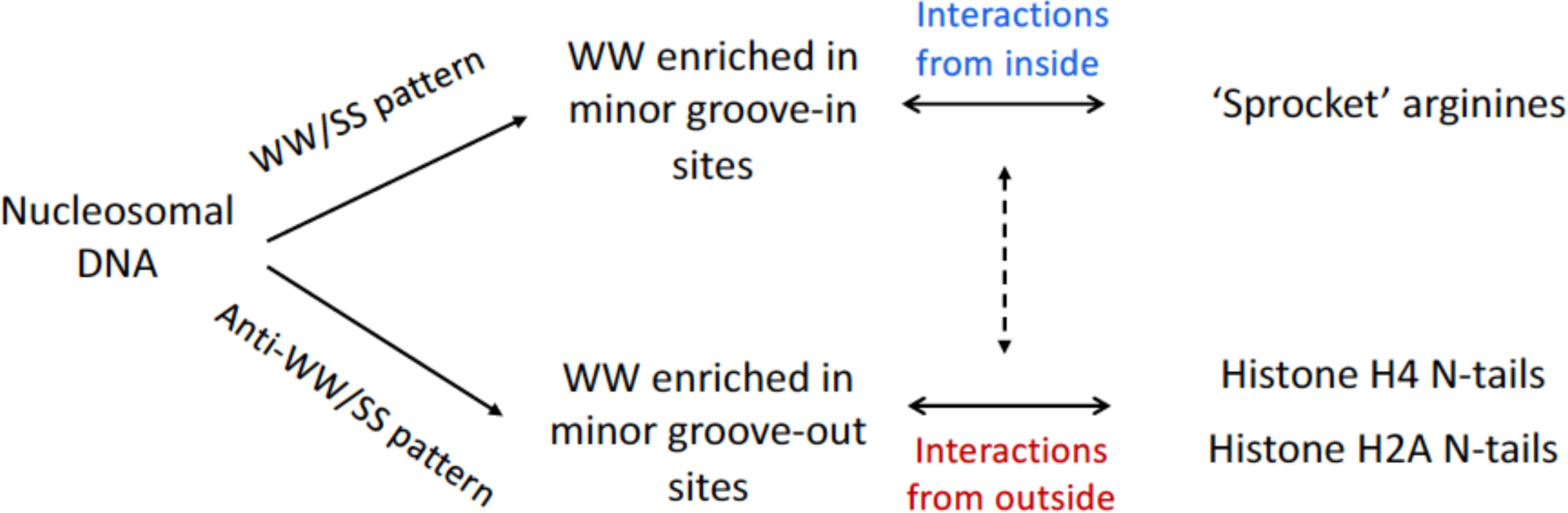
Schematic representation of two types of histone-DNA interactions. Top: Interactions of the first type stabilize ‘canonical’ nucleosomes with the WW/SS sequence pattern. In this case, the WW dinucleotides are predominant at SHL ±0.5, ±1.5, … ±6.5 where the DNA minor groove is facing in (Figure 1). The ‘sprocket’ arginines penetrate DNA minor groove in these locations (5). Bottom: Interactions of the second type stabilize nucleosomes with the anti-WW/SS sequence pattern. Here, the WW dinucleotides are more frequent at locations where the DNA minor groove is facing out. The lysine residues on the H2A and H4 N-tails favorably interact with the AT-rich DNA at SHL ±4 and SHL ±2/±1, respectively (Supplementary Figure S1). Both types of histone-DNA interactions can co-exist in the same nucleosome (indicated by the dashed arrow).

The second type of interactions involves the H2A and H4 histone N-tails, which are in close contact with the minor groove facing out at SHL ±4 and SHL ±2/±1, respectively (Supplementary Figure S1). Our data indicate that the W-T nucleosomes containing N-tails are enriched with the AT-containing fragments at these sites. This trend is clearly opposite to the WW/SS sequence pattern but is consistent with the anti-WW/SS pattern (Figure 5D). Therefore, we hypothesize that the anti-WW/SS nucleosomes are stabilized (at least partially) by the H2A and H4 histone N-tails which favorably interact with the AT-rich minor groove because they are enriched by lysines and arginines. Schematically, the H2A and H4 N-tails act like the arms embracing nucleosomal DNA from outside, whereas the ‘sprocket’ arginines act like the hooks holding nucleosomal DNA from inside (5).

### Relation to other work

Our study was initiated, in part, by the earlier observations of the ‘rogue’ signal near SHL ±1 by Davey (45) and the A *versus* T strand asymmetry near SHL ±4 by Cole *et al.* (43). Various interpretations of these peculiar effects were considered, including interaction of arginine H3R40 with electronegative groups of A:T base pairs near position #65 (45) and wedge formation by AA:TT dimeric steps near #30 (43). However, neither of these explanations account for the position-dependent preference for A over T and T over A (*e.g*., compare positions #30 and #52, Figure 3). It was noted (43) that “an additional factor” is required to elucidate the A *versus* T strand asymmetry in nucleosomes. Our results suggest that interaction of the histone N-tails with the DNA minor groove is likely to be such a factor.

The dynamic nature of these interactions was clearly demonstrated by extensive molecular dynamics (MD) simulations performed by Shaytan *et al.* (46). The histone tails fluctuate in the vicinity of the DNA surface, with the DNA minor grooves serving as kinetic traps. The H2A and H4 N-tails mostly stay within the limits indicated in Supplementary Figure S1, closely resembling the x-ray structure (29). In particular, the H4 N-tails remain most of the time in the minor grooves close to SHL ±1 or SHL ±2; they do not penetrate the minor groove at SHL 0 (near the dyad), at least during the 1-microsecond simulations. The 4-microsecond MD simulations performed later (47) gave essentially the same result – namely, the H4 N-tails don’t go beyond the limits established by the x-ray structure (29). Finally, the very recent NMR study (48) accompanied by 5-microsecond MD simulations has also localized the H4 tails in the DNA region from SHL ±1 to SHL ±2.5.

Note, however, that the H4 N-tail is sufficiently long to penetrate the minor groove close to SHL 0 (Supplementary. Figure S1A). Our result on the high ΔW value in the vicinity of the dyad (Figure 5B) suggests that the H4 N-tail might interact directly with DNA at SHL 0 (but this is not the only possible interpretation). Therefore, it would be interesting to see if the additional 10-fold increase in the time of MD simulations will allow observing the larger-scale fluctuations of the H4 N-tail.

Fan *et al.* (49) analyzed positioning of nucleosomes containing histone H2A.Z on the sea urchin 5S RNA gene. They found that substitution of histone H2A by H2A.Z results in a shi∼ of the predominant nucleosome position by 20 bp, which is the same shi∼ as observed here for the H2A N-tailless nucleosomes (Supplementary Figure S2). This similarity confirms our conclusion on the role played by the histone H2A N-tail in selection of the translational nucleosome positioning, while maintaining the same rotational setting. Interestingly, the same 20-bp shi∼ was observed by Yang *et al.* (50) who analyzed positioning of tailless nucleosomes on the *Xenopus borealis* somatic-type 5S RNA gene. Formally speaking, it would be incorrect to compare directly this 20-bp shi∼ with our result, because in this case all histone tails were digested by trypsin (not only the H2A N-tail). Nevertheless, we can safely summarize that removal of the histone tails can lead to noticeable alteration of the sequence-dependent nucleosome positioning *in vitro*.

## Conclusion

Analyzing distribution of the AT-rich fragments in nucleosomal DNA sequences *in vivo*, we observed non-canonical peaks in the vicinity of superhelical locations SHL ±4 and SHL ±1. These peaks are out of phase with the regularly spaced canonical WW peaks defining rotational positioning of nucleosomes. In the nucleosome X-ray structure (29), DNA at SHL ±4 is in contact with the H2A N-tails, whereas DNA at SHL ±1/±2 is in contact with the H4 N-tails. Therefore, we hypothesized that the observed irregularities in DNA sequence are associated with the favorable interactions between the AT-rich DNA and the lysine-rich histone N-tails.

To test this hypothesis, we reconstituted the H2A- and H4-tailless NCPs on human BRCA1 gene *in vitro* and analyzed the local changes in AT content of these nucleosomes compared to the W-T nucleosomes. We found that the differences between sequences of H2A-tailless and W-T nucleosomes are localized around SHL ±4, while the most pronounced changes in the H4-tailless NCP sequences occur in the central part of a nucleosome (*i.e*., near dyad and at SHL ±1/±2). These results are consistent with the above assumption that the H2A and H4 N-tails are involved in selection of the optimal NCP sequences at SHL ±4 and SHL ±1/±2, respectively. As to the putative interactions between the H4 N-tail and DNA in the vicinity of the dyad (at SHL 0), further structural studies are needed to clarify this question.

Furthermore, we obtained direct evidence for the distinctive repositioning of nucleosomes induced by truncation of the histone N-tails. Importantly, the H2A-tailless nucleosomes remain in phase with the W-T nucleosomes, whereas a significant number of H4-tailless nucleosomes are out of phase with the W-T nucleosomes, suggesting that the N-tails of H2A and H4 histones have different impact on the nucleosome positioning.

The strongest differences between the AT content of the H4-tailless and the W-T nucleosomal sequences are found at SHL 0/±1/±2, where the DNA minor groove faces out (Figure 1). Overall, these locations are enriched with G:C pairs (in accordance with the WW/SS rule), but the H4 N-tails demonstrate partially ‘compensatory’ effect − an increased local affinity to AT-rich DNA (Figure 6). In our interpretation, the H4 N-tail-DNA interactions represent an additional set of histone-DNA contacts complementary to the well-known contacts between ‘sprocket’ arginines and the DNA minor groove. These N-tail-DNA interactions may provide a novel mechanism for stabilization (at least, partial) of the anti-WW/SS nucleosomes that account for ∼25 % of all nucleosomes in the higher eukaryotes (26).

While the structural details of sequence readout by histone tails remain unclear, our data strongly suggest that histone tails preferentially select certain nucleosome positions over other positions, which may have important implications for epigenetic modulation of nucleosome positioning. Certain epigenetic marks, such as acetylation of lysines on histone N-tails, decrease affinity of the tails to the DNA minor groove. This, in turn, may change nucleosome positions and even shift the equilibrium between different chromatin conformation states. Obviously, more studies are needed to elucidate the role of epigenetic marks in positioning of nucleosomes.

## Acknowledgments.

We are grateful to Vladimir Larionov for generous gift of the BRCA1-TAR/YAC/BAC plasmid, as well as to Adam Woolfe and Davood Norouzi for their help at the initial stages of this research.

## Funding

This work was supported by the National Institutes of Health (grant R15GM149587-01 to F.C.) and Intramural Research Program of the NIH, National Cancer Institute (V.B.Z.).

## Supplementary Figures

**Supplementary Figure S1.**
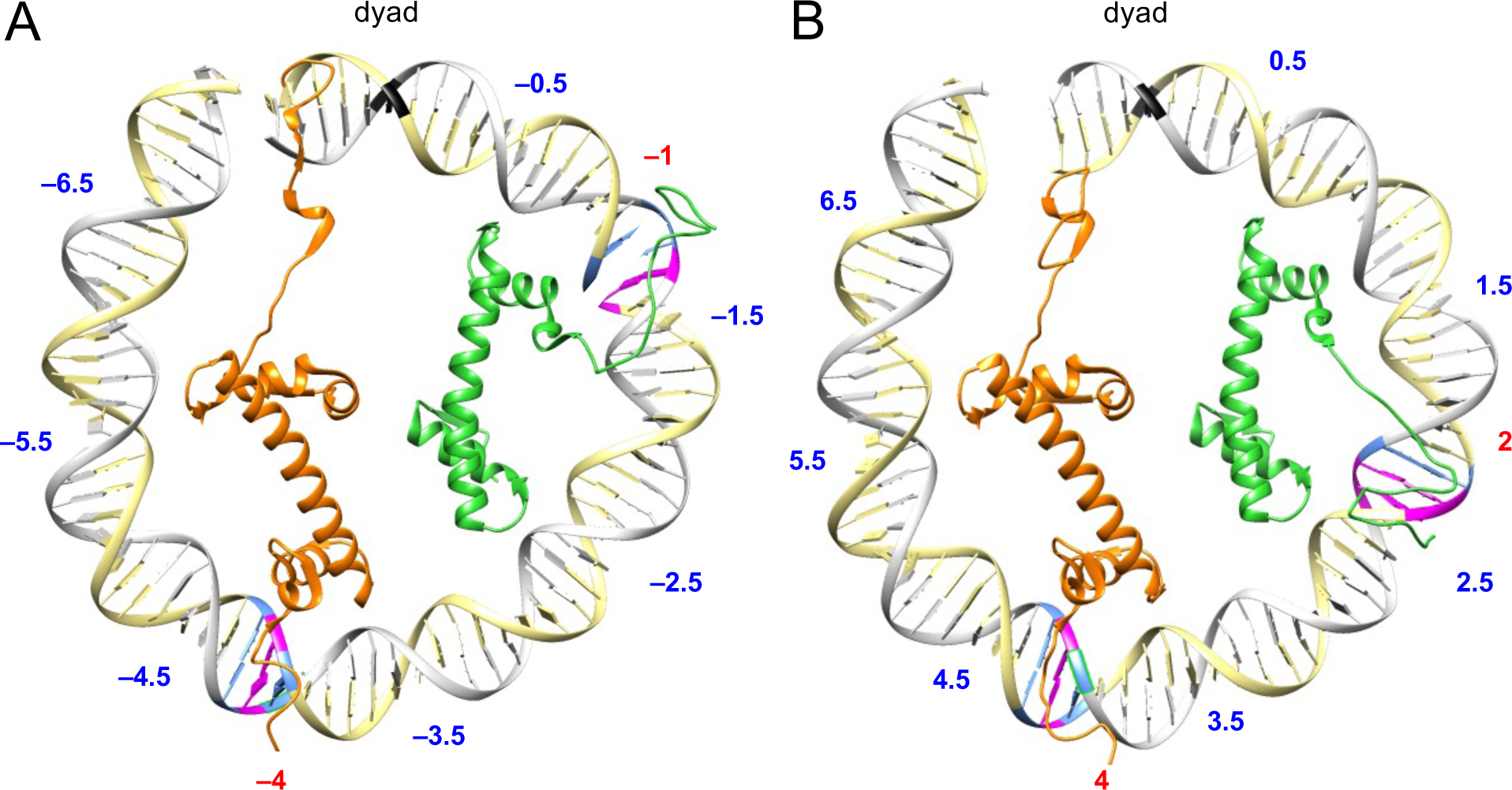
Histone H2A and H4 N-tails in the DNA minor grooves. The ventral (A) and dorsal (B) halves of the nucleosome X-ray crystal structure (29) are shown schematically: the DNA is given in ribbon representation with the dyad colored in black. Only histones H2A (orange) and H4 (green) are shown, whereas the other histones are hidden for clarity. The superhelical locations of DNA (SHL) are numbered from -6.5 to -0.5 in (A) and from 0.5 to 6.5 in (B). The H2A N-tails interact with the DNA minor grooves close to SHL -4/4, while the H4 N-tails penetrate minor grooves between SHL -1 and -1.5 in (A) and between SHL 2 and 2.5 in (B). The nucleotides in close vicinity of H2A and H4 N-tails are colored in blue (A:T pairs) and in magenta (G:C pairs).

**Supplementary Figure S2.**
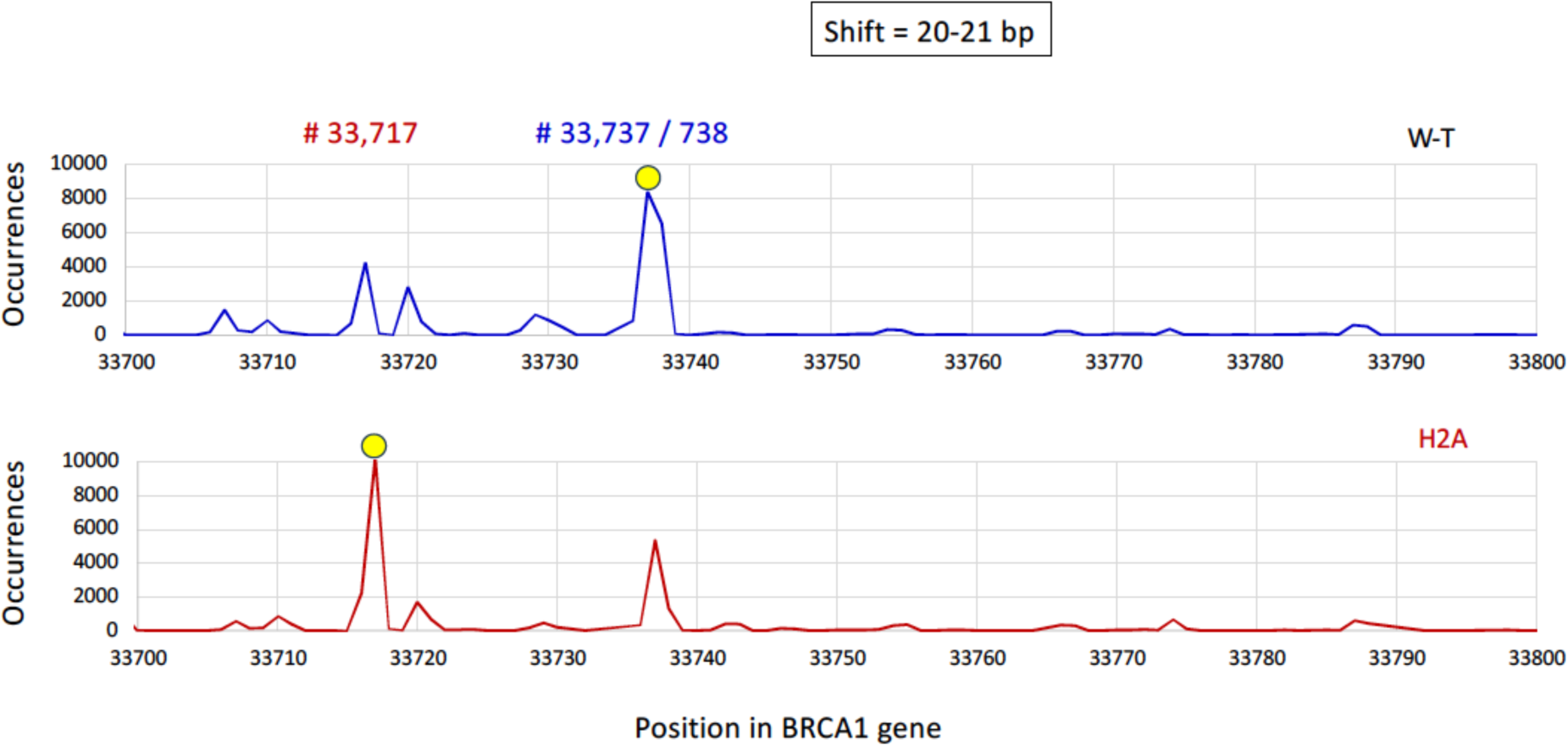
A representative region of BRCA1 gene where the nucleosome occurrences have two strong peaks separated by 20-21 bp. The dominant NCP positions in the W-T and H2A populations of nucleosomes are shifted by 20-21 bp one from the other, so that the NCP positions are in the same phase.

**Supplementary Figure S3.**
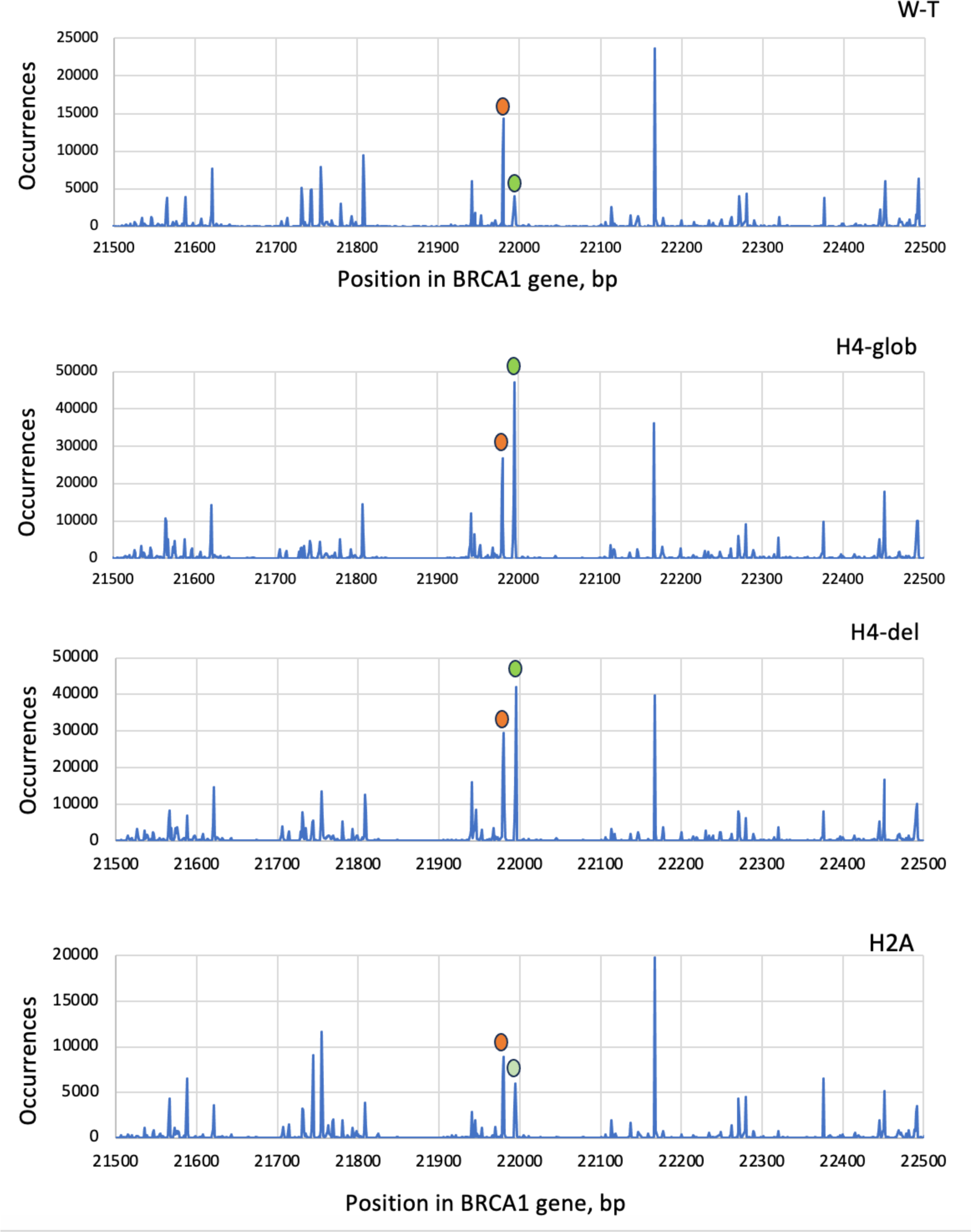
A representative region of BRCA1 gene where the nucleosome occurrences have two strong peaks separated by 15 bp. The nucleosome occurrences in the region 21,500 - 22,500 bp of BRCA1 gene are presented for the four NCP subsets (W-T, H4-glob, H4-del and H2A). Note that in the W-T and H2A subsets, the nucleosomes shown in red represent the dominant NCP positions, whereas in the H4-glob and H4-del subsets, these are the positions shown in green that are dominant. Intriguingly, in the region 21,500 - 22,200 bp there are four peak clusters separated by 190-200 bp, which is remarkably close to the nucleosome spacing *in vivo*. This periodicity does not hold for the whole BRCA1 gene, however (data not shown).

## Notes

### Competing Interest Statement

The authors have declared no competing interest.

