## Supplementary. Figs 1-3 for "Histone N-tails modulate sequence-specific positioning of nucleosomes"

### Supplementary Figures

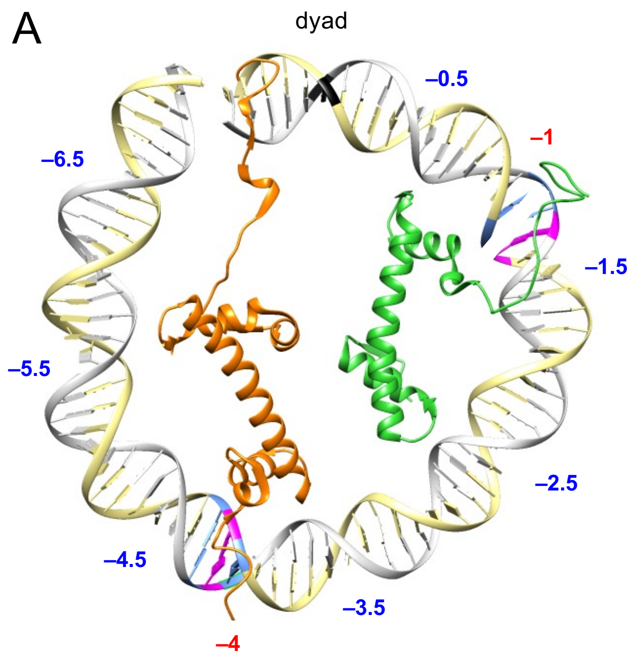


**Supplementary Figure S1.** Histone H2A and H4 N-tails in the DNA minor grooves.

The ventral (A) and dorsal (B) halves of the nucleosome X-ray crystal structure (29) are shown schematically: the DNA is given in ribbon representation with the dyad colored in black. Only histones H2A (orange) and H4 (green) are shown, whereas the other histones are hidden for clarity. The superhelical locations of DNA (SHL) are numbered from -6.5 to -0.5 in (A) and from 0.5 to 6.5 in (B). The H2A N-tails interact with the DNA minor grooves close to SHL -4/4, while the H4 N-tails penetrate minor grooves between SHL -1 and -1.5 in (A) and between SHL 2 and 2.5 in (B). The nucleotides in close vicinity of H2A and H4 N-tails are colored in blue (A:T pairs) and in magenta (G:C pairs).

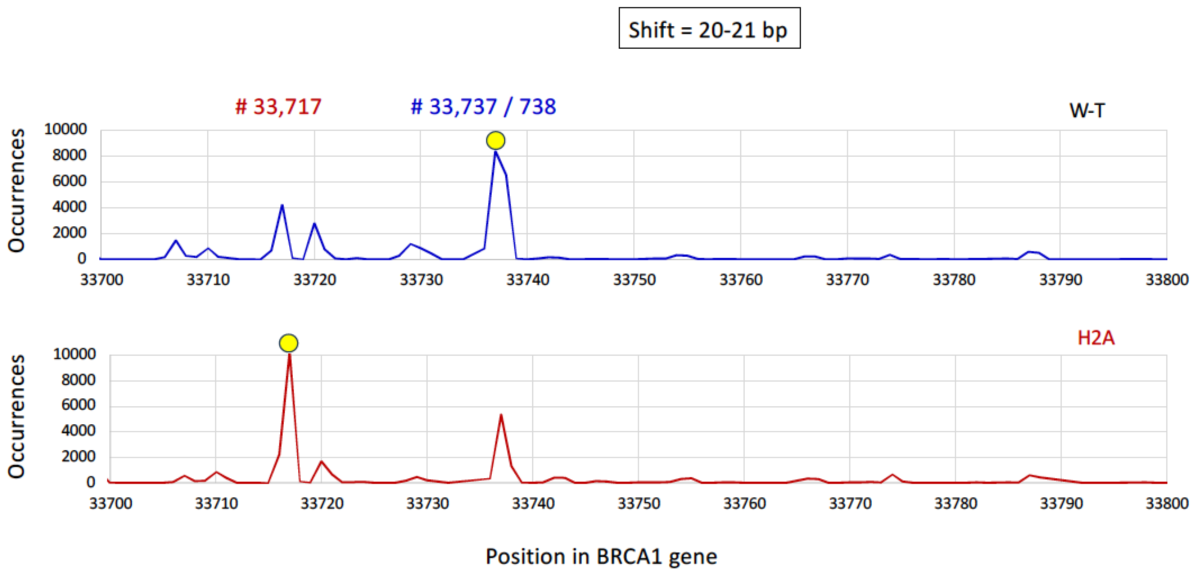


**Supplementary Figure S2.** A representative region of BRCA1 gene where the nucleosome occurrences have two strong peaks separated by 20-21 bp.

The dominant NCP positions in the W-T and H2A populations of nucleosomes are shifted by 20-21 bp one from the other, so that the NCP positions are in the same phase.

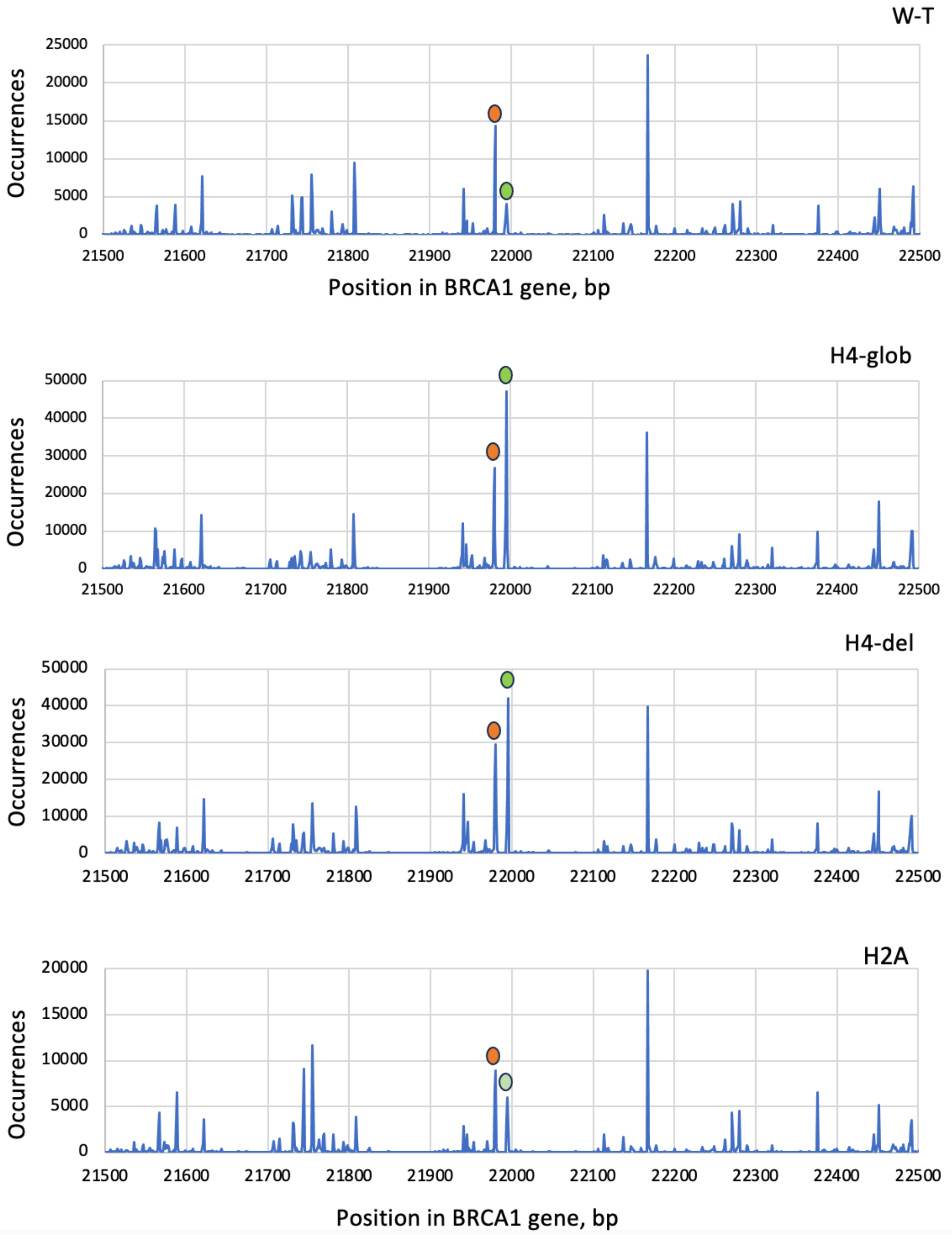


**Supplementary Figure S3.** A representative region of BRCA1 gene where the nucleosome occurrences have two strong peaks separated by 15 bp.

The nucleosome occurrences in the region 21,500 - 22,500 bp of BRCA1 gene are presented for the four NCP subsets (W-T, H4-glob, H4-del and H2A). Note that in the W-T and H2A subsets, the nucleosomes shown in red represent the dominant NCP positions, whereas in the H4-glob and H4-del subsets, these are the positions shown in green that are dominant. Intriguingly, in the region 21,500 - 22,200 bp there are four peak clusters separated by 190-200 bp, which is remarkably close to the nucleosome spacing *in vivo*. This periodicity does not hold for the whole BRCA1 gene, however (data not shown).
